# Populations of genetic circuits are unable to find the fittest solution in a multilevel genotype-phenotype map

**DOI:** 10.1101/817908

**Authors:** Pablo Catalán, Susanna Manrubia, José A. Cuesta

## Abstract

The evolution of gene regulatory networks (GRNs) is of great relevance for both evolutionary and synthetic biology. Understanding the relationship between GRN structure and its function can allow us to understand the selective pressures that have shaped a given circuit. This is especially relevant when considering spatiotemporal expression patterns, where GRN models have been shown to be extremely robust and evolvable. However, previous models that studied GRN evolution did not include the evolution of protein and genetic elements that underlie GRN architecture. Here we use `toy`LIFE, a multilevel genotype-phenotype map, to show that not all GRNs are equally likely in genotype space and that evolution is biased to find the most common GRNs. `toy`LIFE rules create Boolean GRNs that, embedded in a one-dimensional tissue, develop a variety of spatiotemporal gene expression patterns. Populations of `toy`LIFE organisms choose the most common GRN out of a set of equally fit alternatives and, most importantly, fail to find a target pattern when it is very rare in genotype space. Indeed, we show that the probability of finding the fittest phenotype increases dramatically with its abundance in genotype space. This phenotypic bias represents a mechanism that can prevent the fixation in the population of the fittest phenotype, one that is inherent to the structure of genotype space and the genotype-phenotype map.

## Introduction

The evolution of gene regulatory networks (GRNs) is a topic of great relevance [1, 2]. Organisms show a plethora of complex regulatory architectures in order to carry out several developmental programs [3] and to integrate signals from the environment [4]. As a result, much work has been devoted to understanding how these architectures have evolved, and to disentangling the relationship between the structure of a GRN and its function [5, 6]. The underlying motivation is to understand which regulatory motifs appear as a result of selection for a given function or, conversely, what kind of functionality is attained when the structure of the GRNs is determined by other factors. GRNs are also the object of intense research from the standpoint of synthetic biology, which tries to design circuits to perform pre-defined functions [7].

One of regulation’s most interesting outcomes is the generation of spatiotemporal patterns of gene expression that multi-cellular organisms use in their development [8]. Recent work has been devoted to the study of the architecture of GRNs that give rise to different patterns, exploring their robustness and evolvability [9–12]. These studies have found that GRNs can easily evolve to generate new patterns, facilitating the emergence of new developmental programs. The same pattern can be achieved by means of very different mechanisms [11], which in turn determine the levels of robustness and evolvability of the pattern. However, GRNs are the result of interactions between proteins and genetic elements, and the evolution of GRNs is a direct result of changes in protein folding, binding affinities and promoter or enhancer regions. Due to its enormous complexity, models of GRN evolution rarely incorporate these underlying dynamics, although there are some exceptions [13, 14].

Here we use a multilevel computational model of gene regulation to show that some GRN architectures are easier to build from interacting proteins and genes than others. As a result, there is a phenotypic bias [15–17] that turns some GRNs into attractors of evolutionary dynamics, even in the absence of fitness differences.

We focus on Boolean GRNs, in which genes can either be ON or OFF [18, 19]. Although far from other models of gene expression, where the concentration of proteins can vary continuously [20], Boolean networks have been repeatedly used to model GRN evolution [21, 22], and some regulatory functions have been explained best by using Boolean functions [23]. Our Boolean GRNs are also modelled in discrete time, so that the expression of one cell in time *t* + 1 is determined by its expression and that of its neighbouring cells in time *t*. This formalism transforms GRNs into cellular automata [24]. Connecting several cells in a one-dimensional tissue, and allowing for propagation of gene products between neighbouring cells, we obtain spatiotemporal patterns that are similar to those found in real organisms.

These Boolean GRNs are built on top of a simple model of cellular biology, `toy`LIFE [25,26]. `toy`LIFE organisms contain genes, which are translated into proteins that interact with each other to form dimers. Both dimers and proteins alter the expression of genes, thus creating Boolean GRNs like the ones described above. As a consequence, `toy`LIFE is a multilevel map from binary genomes (genotypes) to Boolean GRNs (first phenotype level) to cellular automata (second phenotype level) to spatiotemporal patterns (third phenotype level) (Figure 1), thus allowing us to study the effects of molecular evolution at different phenotypic levels.

**Figure 1:**
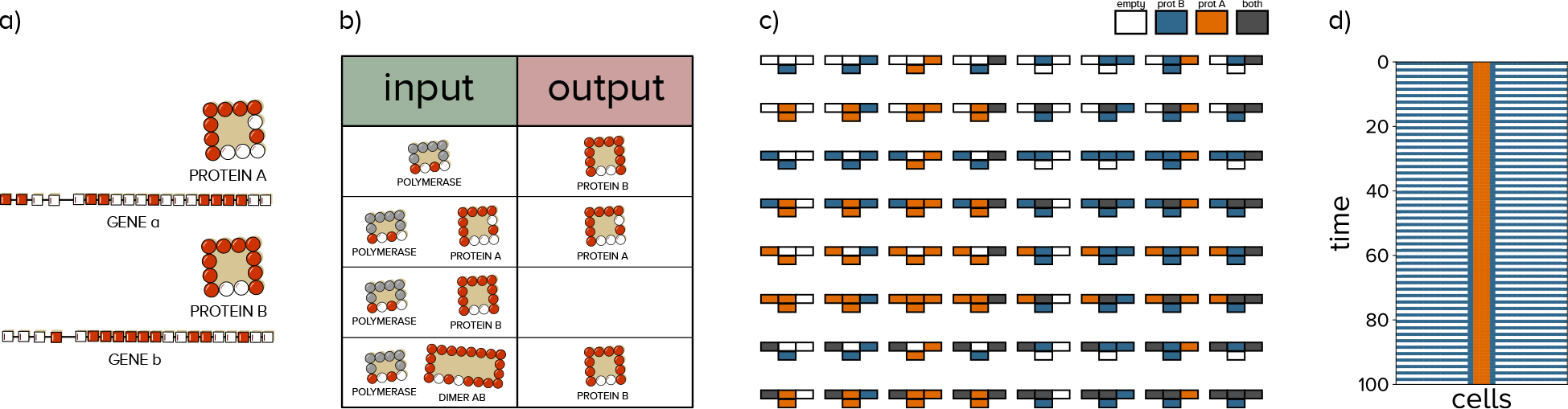
toyLIFE is a multilevel genotype-phenotype map. **(a)** `toy`LIFE genotypes are binary strings of length 20*n*, where *n* is the number of genes in the genome. The first 4 letters of each gene represent its promoter region, while the remaining 16 are the coding region. The coding region, when expressed, turns into a protein that folds into a 4 × 4 lattice (see Supplementary Text Section 1). **(b)** Following `toy`LIFE’s interaction rules, we obtain the corresponding gene regulatory network (GRN), represented here by its truth table. **(c)** Each GRN determines, under some propagation rules, a unique cellular automaton. Given the state of a cell and its neighbours at time *t*, `toy`LIFE’s rules determine the state of the cell at time *t* + 1, where cells can be empty (white), expressing protein *A* (orange), expressing protein *B* (blue) and expressing both proteins (grey). **(d)** Under certain initial conditions (in this case, the expression of protein *A* in the middle cell of the tissue), the cellular automata give rise to spatiotemporal patterns of gene expression. In this case, the cellular automaton in **c** leads to an alternating pattern in which the tissue expresses protein *B* and then doesn’t express anything, while in the centre of the tissue three cells express protein *A* continuously.

We show that `toy`LIFE genomes with two genes are able to generate a wide variety of GRNs and spatiotemporal patterns. Moreover, not all of these are equally abundant in genotype space: some GRNs are mapped by many genotypes, while others are comparatively rare. We find that this phenotypic bias is enough to steer evolving populations towards more abundant GRNs, thus introducing an additional element when trying to explain GRN evolution, one that is not related to function or structure. Furthermore, we also show that this phenomenon can result in the inability of the evolutionary search to find some regulatory patterns, even when they are fitter than every other.

## Results

### Boolean networks and spatiotemporal patterns

In a Boolean GRN, a gene can either be ON (expressed) or OFF (not expressed). The expression state of each gene at time *t* + 1 is a function of the expression states of all the other genes in the network at time *t*, so that each state of the network maps into another state. In a GRN with two genes *a* and *b* and corresponding proteins *A* and *B*, this mapping can be represented as follows:

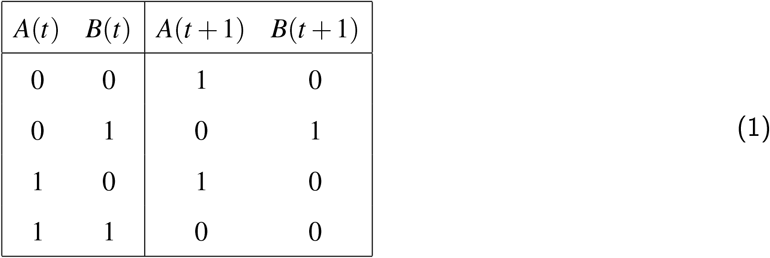

This representation is called a truth table, connecting every input state to an output state. In this case, the state (0, 0) is mapped to (1, 0), which means that gene *a* is expressed constitutively. The next two rows indicate that both *a* and *b* activate their own expression, while the last row shows that both genes repress each other. The truth table determines the temporal expression patterns of a Boolean GRN, thus giving us all the information we need to study this system.

We want to study the spatiotemporal patterns of two-gene GRNs embedded in a one-dimensional tissue. First, we define the number of cells in the tissue, which we will consider to be constant. For our purposes, we choose tissues with 31 cells in a row. The number of cells is arbitrary and it does not affect our results: the same patterns are generated by the same truth tables under similar regulatory inputs (Supplementary Figure S8), so no phenomenology is lost from restraining our study to this tissue size.

Now we define the connections between different cells in the tissue. We will assume that only protein *A* can propagate to the adjoining cells (Figure 2). As a result, the input state of cell *c*_*i*_ in time *t* + 1 will be affected by the output states of cells *c*_*i*−1_ and *c*_*i*+1_ in time *t* —as well as its own. We will further assume that there is enough protein *A* to stay inside the cell and propagate to the adjoining ones. For the cells at the beginning and end of the tissue, we impose the following boundary condition: cell *c*_0_ will be affected by itself and cell *c*_1_, and cell *c*_*L*_ will be affected by itself and *c*_*L*−1_ —remember that *L* = 30 throughout.

**Figure 2:**
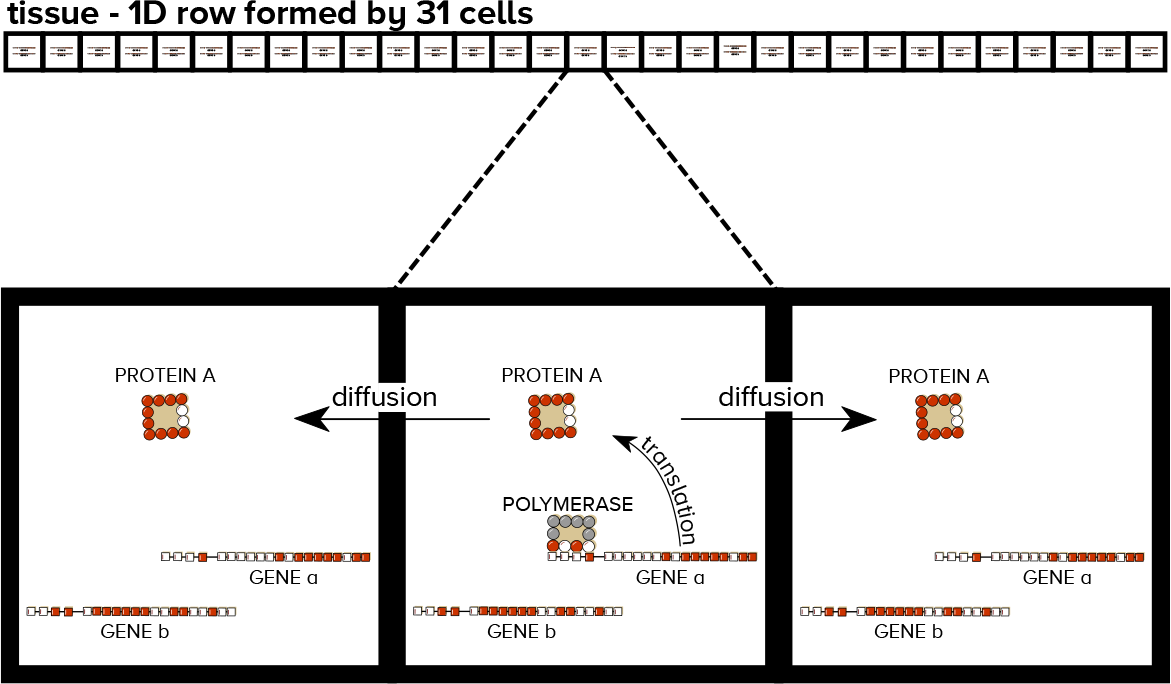
Pattern-formation phenotype in toyLIFE. We consider a one-dimensional row formed by 31 cells. The figure illustrates one example of how this multicellular phenotype works using toyLIFE for illustration purposes. When protein *A* is expressed, it can propagate to neighbouring cells and influence gene expression there. This way, the spatiotemporal state of the tissue becomes a cellular automaton.

With these rules, each GRN (defined by a truth table) gives rise to a cellular automaton [24] (Figure 1c) with four states: (0) no protein is expressed (white), (1) protein *B* is expressed (blue), (2) protein *A* is expressed (orange) and (3) both proteins are expressed (grey). Cellular automata are compactly described by the output they produce given an input. Because the input of a cell is formed by itself and its adjoining cells, and because each of them can be in 4 states, the number of input states is 4^3^ = 64. The number of possible cellular automata is, therefore, 4^64^ ≈ 3.4 × 10^38^. We will see below that the number of two-gene `toy`LIFE genotypes, which we use to generate our GRNs, is around 10^12^, not enough to explore this vast number.

Each cellular automaton, in turn, gives rise to a spatiotemporal pattern that will depend on the initial conditions of the tissue at time *t* = 0 and the external input received. It is soon evident that the space of these scenarios is hyper-astronomical in size [17], and so we choose to start our dynamics with both genes in every cell in the OFF state, except the cell in the middle of the tissue (*c*_15_), where we will express protein *A*, modelling a signal received from the exterior of the tissue. We then explore the expression dynamics of the whole tissue for 100 time steps, enough to resolve all patterns.

We now explore four relevant GRNs and their resulting patterns (Figure 3). They are (a) a double-negative feedback loop with self-activation, (b) the same as before but with gene *a* having constitutive expression, (c) a double-positive feedback loop with self-activation and (d) a double-positive feedback loop without self-activation. Figure 3 shows the truth tables associated with these GRNs and the patterns they generate under the conditions mentioned above. The first two patterns result in protein *A* being expressed in a stable manner in the whole tissue. The difference between them is that in Figure 3b protein *A* is expressed constitutively in every cell, while in Figure 3a that signal must propagate through the tissue. The pattern in Figure 3c is similar to the one in Figure 3a, but both proteins end up expressed in the tissue, as a result of the positive feedback loop. Finally, in Figure 3d the tissue expresses protein *A* and *B* in an alternating way.

**Figure 3:**
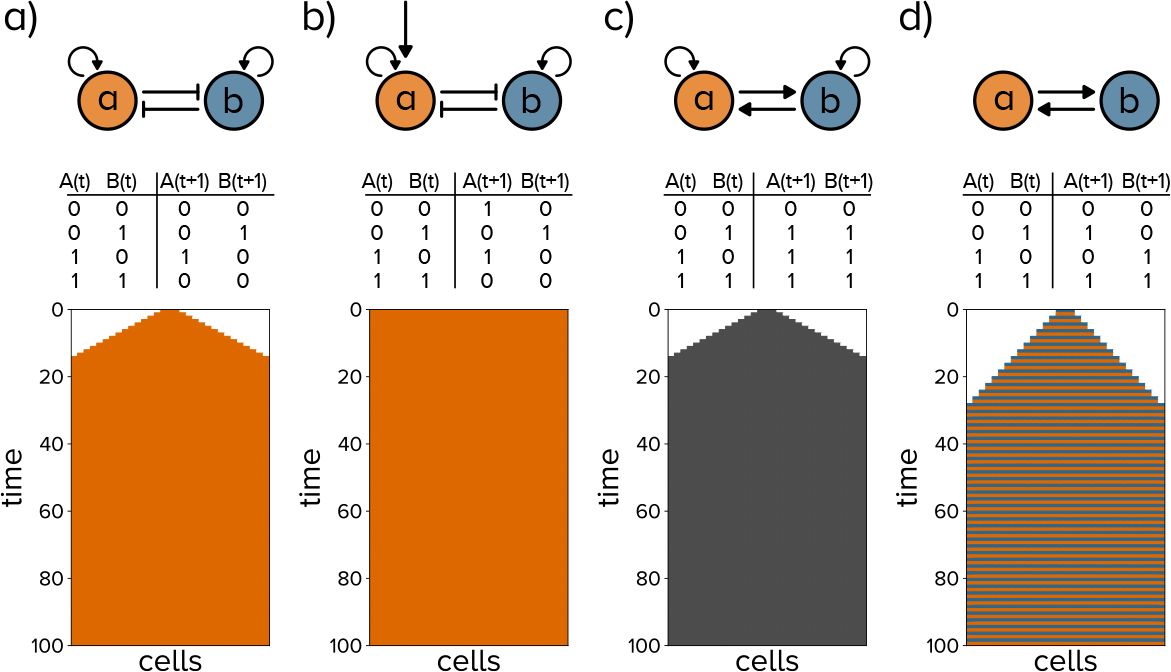
Some examples of patterns generated by two-gene Boolean GRNs. **(a)** A double-negative feedback loop, with self-activation, results in a pattern that expresses protein *A* (orange) stably and expanding through the tissue. **(b)** A double-negative feedback loop with self-activation, where gene *a* is expressed constitutively, leads to the whole pattern expressing protein *A* stably through time. **(c)** A double-positive feedback loop with self-activation loops leads to both proteins *A* and *B* (grey) being expressed in the tissue in a stable way, and expanding through the tissue. **(d)** A double-positive feedback loop without self-activation leads to an alternating pattern where the tissue expresses first protein *A* (orange), then protein *B* (blue), and so on. Notice how the speed with which the pattern extends throughout the tissue is half the speed of patterns in **a** and **b**. This is because only protein *A* is allowed to propagate to the neighbouring cells (see Figure 2), so that the pattern can only extend when protein *A* is expressed.

Let us focus on the pattern generated by the network in Figure 3b. There are sixteen GRNs that generate the same pattern under the conditions defined above (Supplementary Figure S9 shows the truth tables for all of these). If there were selection pressures to create that particular pattern, we could expect evolutionary dynamics to choose among these sixteen GRNs with equal probability, everything else being equal. This is certainly what almost every mathematical model of phenotypic evolution (including previous models of GRN evolution) would predict.

We performed Wright-Fisher evolutionary simulations with `toy`LIFE organisms in a strong selection, weak mutation regime (Methods), and selected the pattern in Figure 3b as the evolutionary target —i.e. we assigned maximal fitness to it, and every other pattern became less fit as it differed more from the target (see Methods for the complete definition of the fitness function). We found that, after 100, 000 mutations, 93% of simulations ended up finding one particular GRN among all sixteen (GRN XI in Figure S9, see below), and the network in Figure 3b (GRN V) does not appear as the endpoint of evolutionary dynamics in any of the 1, 000 simulations. In order to understand this somewhat unexpected result, we now discuss how Boolean GRNs are obtained from `toy`LIFE genotypes.

### Regulation in toyLIFE

We will introduce gene regulation in `toy`LIFE through an example (for an in-depth discussion of `toy`LIFE’s rules, see Supplementary Text Section 1 and [26]). Consider the genotype in Figure 4a. Proteins *A* and *B*, the expression products of genes *a* and *b*, respectively, bind together to form dimer *AB* (Figure 4b). Due to `toy`LIFE’s interaction rules, the expression of gene *a* is activated by protein *A*, its own expression product. On the other hand, the expression of gene *b* is activated by the polymerase (it is a constitutively expressed gene), but it is inhibited by both proteins *A* and *B*. The dimer does not bind to any promoter (Figure 4c). With this information, we can compute the expression output of this genotype given each input, i.e. its truth table (Figure 4d). When no protein is present, the polymerase (which is always present in the cell) will activate gene *b* and the output will consist of protein *B*. The same will happen if dimer *AB* is present in the cell: because it does not interact with either promoter, the polymerase will activate the expression of gene *b* again. If protein *A* is present, it will displace the polymerase and gene *b* will not be expressed, but *A* will also activate its own expression. Finally, if protein *B* is present, it will inhibit its own expression, and nothing will be expressed in the cell. In this way, we map a binary sequence (coding for the genome’s two genes) into a Boolean GRN. It is interesting to note that this regulatory function cannot be expressed with an arrow diagram similar to those in Figure 3: there is no way to represent the overriding effect that the dimer has on each protein’s regulatory logic using this kind of diagram.

**Figure 4:**
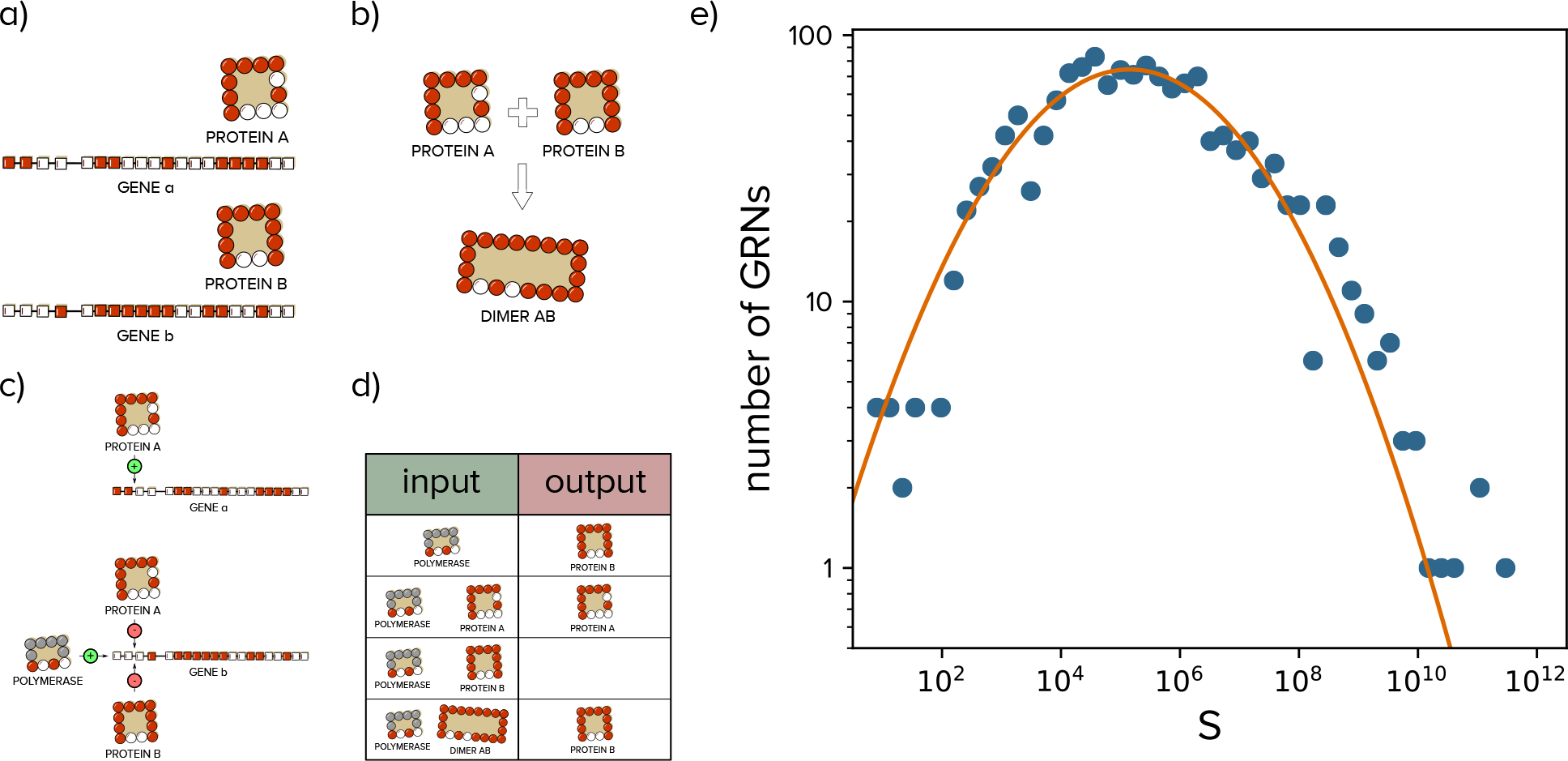
Regulatory logic in toyLIFE. **(a)** Example of a two-gene genotype in toyLIFE. These genes express strings of 16 amino-acids that fold into a 4 × 4 lattice, following the rules of the HP model (Supplementary Text Section 1). **(b)** Protein *A* and protein *B* can bind together to form dimer *AB*. **(c)** Regulatory logic of genes *a* and *b*. Protein *A* inhibits its own expression. The polymerase activates the expression of protein *B*, while both protein *A* and *B* inhibit it. The dimer *AB* does not bind either promoter. **(d)** Truth table representing the regulatory logic of this two-gene genotype, obtained from the information in **c**. See text for details. **(e)** Not all GRNs generated by `toy`LIFE two-gene genotypes are equally likely in genotype space. In fact, the distribution of abundances (*S*) follows a log-normal distribution (*R*^2^ = 0.94).

The cellular automaton is now uniquely determined by the GRN once we take into account two additional input states: protein *A* plus dimer, and protein *B* plus dimer. These two states can appear as a result of protein products propagating from one cell to the next. With this information we can unequivocally compute each genotype’s corresponding cellular automaton.

It is worth noting that in the process of defining these phenotypic levels we have already introduced a lot of degeneracy. For instance, there are 2^40^ ≈ 10^12^ genotypes with two genes, but they only give rise to 1, 472 different GRNs, which in turn generate only 453 different cellular automata —an average of ≈ 2 × 10^9^ genotypes per cellular automata. Not all GRNs are equally probable in genotype space, however: the distribution of abundances of GRNs follows a log-normal distribution (Figure 4e), which has been observed in many other genotype-phenotype models and has been shown to be universal under some very general assumptions [16, 26–28]. The most abundant GRN is mapped by more than 500 billion genotypes, while the rarest one is only mapped by 8 genotypes. This phenomenon has been called phenotypic bias [15, 16], and it is also observed in the distribution of abundances of cellular automata (Supplementary Figure S10a). As a consequence, the sixteen GRNs that generate the pattern in Figure 3b (Supplementary Figure S9) also have varying abundances. The most common one is GRN XI, mapped by 2.017 × 10^11^ genotypes, roughly 18% of all two-gene genotypes in `toy`LIFE. Its truth table is

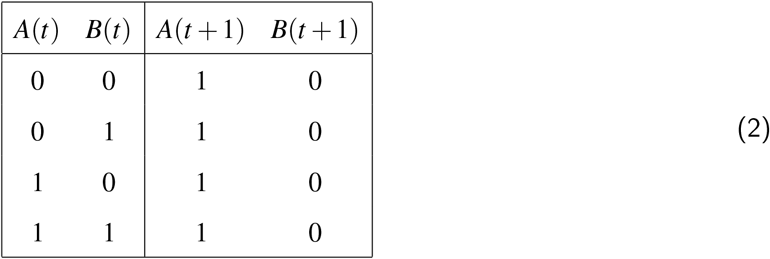

This is, admittedly, a very simple Boolean GRN, in which every input state leads to the same output: that of protein *A* being expressed. Previous work [29] has argued that simpler phenotypes should be more abundant in genotype space, and this is indeed what we observe at all phenotypic levels (Supplementary Figure S11). In comparison, the least abundant Boolean GRN among these sixteen (GRN VIII) is mapped by just 203, 641 genotypes, a million times less abundant than GRN XI. Finally, the double-negative feedback loop in Figure 3b that we were searching for originally (GRN V) is mapped by 9.4 × 10^6^ genotypes, which is 5 × 10^−5^ times less abundant than GRN XI. As a result of this phenotypic bias in the sixteen GRNs, when we evolve populations of `toy`LIFE organisms to express this simple pattern as described above, populations find GRN XI 93% of the times —although all sixteen are equally fit in this scenario. In fact, the proportion of times our simulations end up in a particular GRN closely reflects its relative abundance in genotype space (Supplementary Figure S12). In other words, introducing an additional level to the GRN-to-pattern genotype-phenotype map causes a bias in the abundances of different GRNs, which in turn affects evolutionary dynamics [30].

### Pattern formation in toyLIFE

The differences in abundances in the Boolean GRNs generated by `toy`LIFE are magnified at the pattern level (Supplementary Figure S10b): some patterns are mapped by billions of genotypes, while others are generated by only hundreds of them. This difference is critical, as we will see now. Suppose an evolutionary scenario where we select for pattern 113, shown in Figure 5a. This pattern is mapped by only 5, 312 genotypes, so finding it in genotype space seems hard *a priori*. However, naïıve evolutionary predictions would say that, being the fittest phenotype, it should be eventually selected and fixed in the population. When we perform the evolutionary simulations with pattern 113 as the target (Methods), it appears as the evolutionary endpoint only in 3% of the 1, 000 simulations. Instead, the pattern that appears in most of the simulations is pattern 109 (Figure 5b), which has a fitness of 0.991 relative to that of pattern 113, and is mapped by 1.6 × 10^8^ genotypes. There are 33, 280 mutational paths between pattern 109 and pattern 113 (counted as the number of pairs of genotypes mapping to each pattern that are one point mutation apart), so populations expressing pattern 109 could eventually find pattern 113 without having to go through any fitness valley. However, this number represents only 0.0005% of all connections from pattern 109 to other phenotypes: finding pattern 113 from pattern 109 is truly like finding a needle in a haystack. In other words: the phenotypic bias towards pattern 109 is enough to counteract pattern 113’s fitness benefit. Curiously enough, pattern 170 (Figure 5c), which is not very similar to pattern 113, with a fitness of 0.54, also appears frequently as the endpoint of our simulations. In this case, there are no mutations from pattern 170 to 113, so it seems that some populations quickly find pattern 170 as a suboptimal fitness peak, and then become trapped in it, as there are no mutations to fitter alternatives.

**Figure 5:**
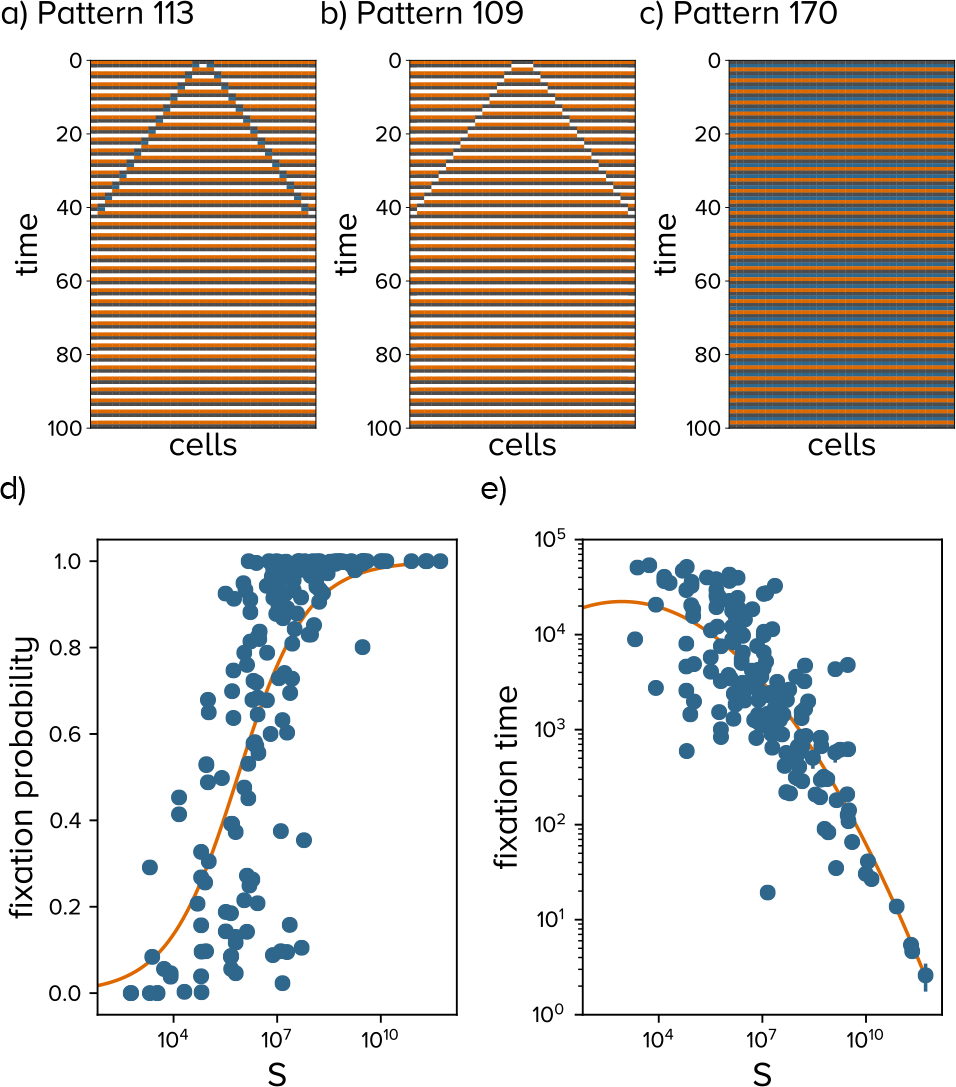
Evolving populations are not able to find rare patterns, even when they are fitter. **a)** Pattern 113 is rarely found in our evolutionary simulations. **b)** Pattern 109 is similar to 113, but it is generated by 1.64 × 108 genotypes —about 105 times commoner. As a result, it appears as the endpoint of our simulations 84% of the times. **c)** Pattern 170 also appears as the endpoint of the simulations 8% of the times, even though it is not very similar to pattern 113. This is due to its high abundance in phenotype space: 1.36 × 108 genotypes are mapped to it. **(d)** This phenomenon is not restricted to pattern 113. The probability of finding a target pattern (*p*) goes to zero as the logarithm of pattern abundance (*S*) decreases. Line: *p* = (1 + (430767/*S*)^1/2^)^−1^, R^2^ = 0.58. **(e)** Even when simulations do find the fittest pattern, the time to reach it (*T*) increases as pattern abundance decreases. Line: log_10_ *T* = 4.35 − 0.05(log_10_ *S* − 2.93)^2^, R^2^ = 0.68.

This result means that some patterns will not be reachable by evolution, not because they are less fit, but because they are very rare in genotype space. This phenomenon is true for every rare pattern, and indeed we see it in simulations where each of the 172 patterns obtained in our system is set as the target of evolution. The probability of finding the fittest pattern decreases dramatically with pattern abundance (Figure 5a). And, even if the pattern is found, the time to find it decreases super-exponentially with pattern abundance (Figure 5b).

## Discussion and Conclusions

The main intention of this work is to show that the complex mapping from DNA sequences to genetic circuits in real cells is in all likelihood biased towards some GRNs, so that some of them are much more common in genotype space. The results of our computational simulations show that this bias is enough to prevent populations from finding the fittest phenotype [15]. Several mechanisms had been previously proposed to explain why populations do not reach the fittest solution, such as frequency-dependent selection [31] or the fittest vs the flattest [32, 33] —which do not apply here, as our populations are always homogeneous. However, phenotypic bias is the first of these mechanisms that arises out of the intrinsic structure of the evolutionary search space, and it is completely independent from population effects and from the structure or function of the GRN being selected for. In this sense, phenotypic bias is playing in evolutionary dynamics the same role that entropy plays in statistical physics. The entropy of a macrostate is related to the number of microstates that are consistent with it without altering the properties that characterise the system. In statistical physics, macrostates are typically described by macroscopical properties such as temperature, pressure or volume, while microstates differ in the positions and velocities of individual particles. In evolutionary dynamics, there is a natural analogy between microstates and genotypes, on the one hand, and macrostates and phenotypes, on the other [34]. The conflict between energy and entropy found in physical systems is the same we have found between fitness and phenotypic bias, and the trapping by abundant phenotypes is akin to a glassy dynamics in physical systems [35].

Our results cannot be explained by phenotypic bias alone, however. In the simulations to find rare pattern 113, we found that a non-negligible fraction of simulations ended in pattern 170, that had a fitness of 0.54 but an abundance in genotype space that was similar to pattern 109, a fitter alternative. The reason populations got stuck in pattern 170 is because genotype space is structured as a complex network, and not all paths from one pattern to the other are actually possible. In this case, there are no connections between pattern 170 and either pattern 109 or 113, so once the population has found this local fitness peak, there is no way it can reach the other, fitter alternatives under our selection regime. The effect of networked genotype spaces on evolutionary dynamics is far from trivial and has yet to be disentangled [36]. Further work has to be devoted to study its effects in this particular system.

The consequences of this work are immediate for the evolution of genetic circuits. Our results suggest that some ideal solutions could be hard to find in genotype space, and that evolution has had to work with more abundant, less efficient alternatives. However, the number of available phenotypes grows very quickly with genotype size in many computational genotype-phenotype maps [17, 26, 27], and so it is reasonable to expect that evolution could always find alternatives that are, if not optimal, at least highly functional. On the other hand, synthetic biologists trying to design a particular circuit could be aiming at a particularly rare structure, which would make its *a priori* evolution very unlikely. This would make that circuit very unstable in evolutionary terms, and mutations could easily change it into a different circuit, with undesired functions.

In relation to this, our results also suggest that phenotypic bias will have an effect on both robustness and evolvability. Previous models studying these properties in GRNs [11] found that they depended on the mechanism by which a GRN generates a pattern. Our results add a new layer, showing that more abundant GRNs will generate more robust patterns, independently on their mechanism or structure. Thus, understanding which GRNs are more abundant in genotype space is essential to unravel the evolution of robustness and evolvability.

We are aware of the limitations of `toy`LIFE as a discrete-time Boolean model to model continuous-time, stochastic protein concentration dynamics. However, phenotypic bias is not a particular characteristic of `toy`LIFE and is rather very common in computational genotype-phenotype maps [15, 16, 29]. Thus, our main results are not limited to this particular choice of model, and they could be easily extended to other, more realistic genotype-phenotype maps. On the other hand, `toy`LIFE is a very convenient model to study multilevel genotype-phenotype relationships [26], which are complex and largely unknown. This model potential to generate complex behaviours is yet to be explored fully.

## Methods and Materials

### Fitness

The fitness function for our evolutionary simulations is calculated as follows: each pattern is a string in base four of length *L* = 31 · 100. For every evolutionary scenario, we choose one particular pattern *p*_T_ as the target value, and assign fitness 1 to it. Then we compute the Hamming distance *D* of a pattern *p* to the target as

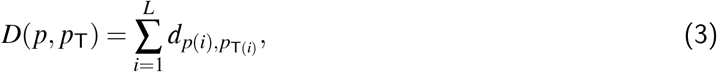

where *d*_*i*, *j*_ is Kronecker’s delta, which is equal to 1 if *i* = *j* and 0 otherwise, and *p*(*i*) is the *i*-th letter in the string *p*. Fitness is then calculated as

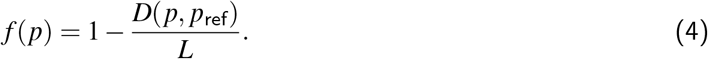

### Evolutionary simulations

We assume a strong selection, weak mutation scenario. In this regime, Wright-Fisher dynamics are reduced to a continuous-time random walk in genotype space. We only consider point mutations, which arise in the population at constant rate *μ*, and the fixation rate of a new mutation is given by

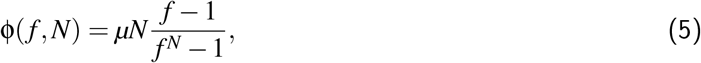

where *f* is the fitness of the current phenotype relative to that of the mutant, and *N* is population size [37]. We assume *μ* = 1, which is equivalent to counting time in mutations instead of generations. Genotypes are binary strings of length 40, which are mapped to a pattern using `toy`LIFE’s rules (Supplementary Material Section 1). We start the simulations choosing a genotype at random, and then simulate populations dynamics using Gillespie’s algorithm [38]. We simulated populations of size *N* = 10, 000 for *T* = 100, 000 mutations, and repeated this process for *R* = 1, 000 replicates for each of the experiments. The choice of population size was made so that deleterious mutations were hardly never accepted.

### Phenotypic complexity

Following [29], we approximate the algorithmic complexity of a binary string *x* = {*x*_1_ … *x*_*n*_} as

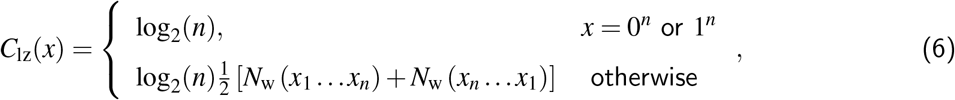

where *n* = |*x*| and *N*_*w*_(*x*) is the number of words in the dictionary created by the Lempel-Ziv algorithm [39]. For each phenotypic level (GRNs, cellular automata and patterns), we translate each base-four string identifying the phenotype to binary code, and then compute *C*_lz_. So, for instance, string 312011 would become 110110000101. GRNs are represented as a binary string by reading all the output entries in the truth table (*GRNII* in Supplementary Figure S9 is equivalent to 10001001) and then adding the output states of the two additional input states mentioned in the main text: protein *A* plus dimer, and protein *B* plus dimer. Therefore, GRNs can be univocally represented as binary strings of length 12. Cellular automata are base-four strings of length 64 that become binary strings of length 128 after converting from base four to base two. Finally, patterns are base-four strings of length 3, 100 that become binary strings of length 6, 200.

## Acknowledgments

PC is supported by a Ramón Areces Postdoctoral Fellowship. This research has been supported by Ministerio de Ciencia, Innovación y Universidades/FEDER (Spain/UE) through grant PGC2018-098186-B-I00 (BASIC) and and FIS2017-89773-P (MiMevo).

## Code availability

`toy`LIFE and all the code used to obtain the results in this paper is freely available at https://github.com/pablocatalan/toylife/.

## Supplementary Text

### 1 toyLIFE

`toy`LIFE was originally presented in [25]. We give here its main details, with slight modifications in the definition of the model, as presented in [26].

#### 1.1 Building blocks: genes, proteins, metabolites

The basic building blocks of `toy`LIFE are toyNucleotides (toyN), toyAminoacids (toyA), and toySugars (toyS). Each block comes in two flavors: hydrophobic (H) or polar (P). Random polymers of basic blocks constitute toyGenes (formed by 20 toyN units), toyProteins (chains of 16 toyA units), and toyMetabolites (sequences of toyS units of arbitrary length). These elements of `toy`LIFE are defined on two-dimensional space (Supplementary Figure S1).

**Supplementary Figure S1:**
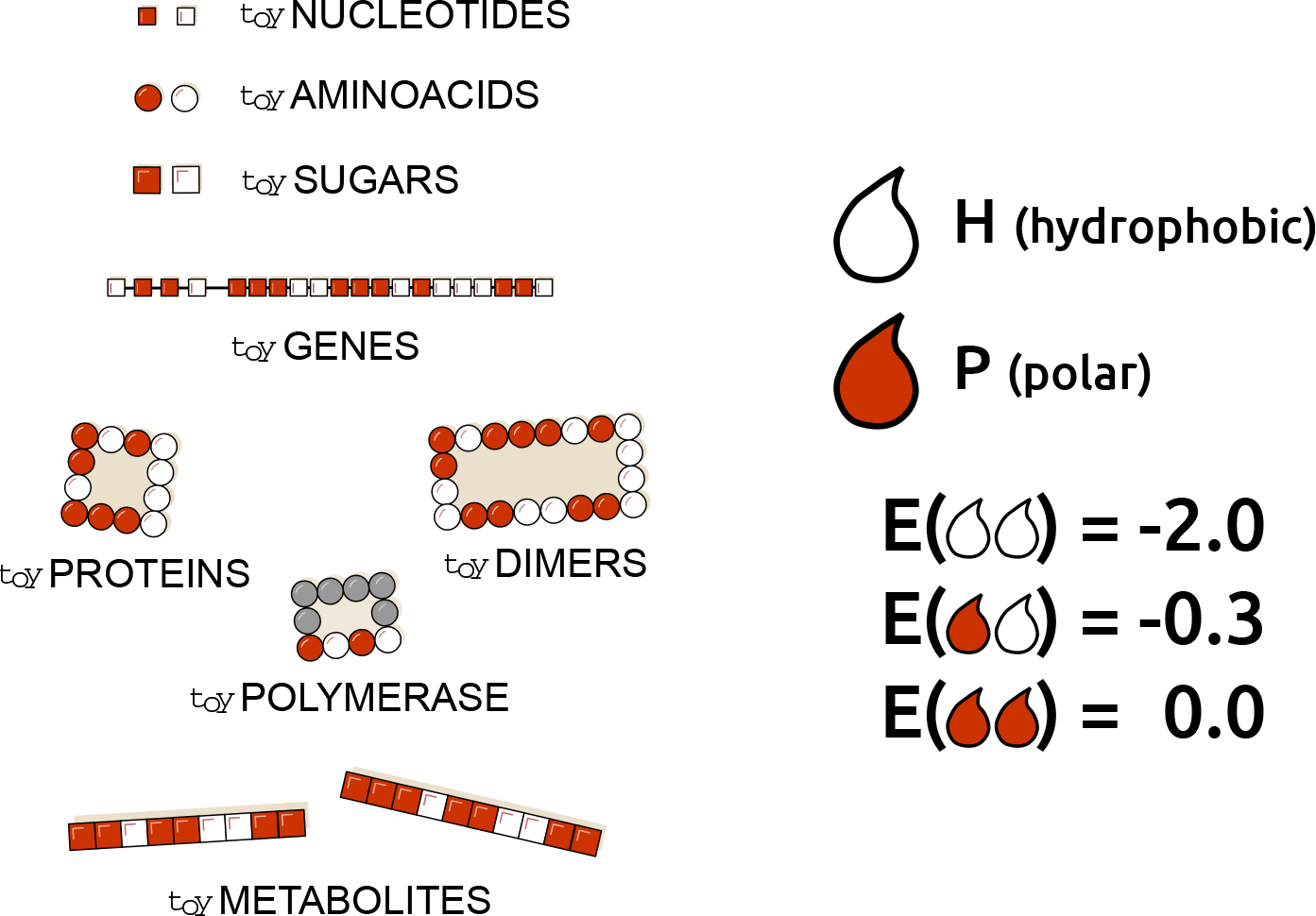
Building blocks and interactions defining toyLIFE. The three basic building blocks of `toy`LIFE are toyNucleotides, toyAminoacids, and toySugars. They can be hydrophobic (H, white) or polar (P, red), and their random polymers constitute toyGenes, toyProteins, and toyMetabolites. The toyPolymerase is a special polymer that will have specific regulatory functions. These polymers will interact between each other following an extension of the HP model (see text), for which we have chosen the interaction energies *E*_HH_ = −2, *E*_HP_ = −0.3 and *E*_PP_ = 0 [40].

##### toyGenes

toyGenes are composed of a 4-toyN promoter region followed by a 16-toyN coding region. There are 2^4^ different promoters and 2^16^ coding regions, leading to 2^20^ ≈ 10^6^ toyGenes. An ensemble of toyGenes forms a genotype. If the toyGene is expressed, it will produce a chain of 16 toyA that represents a toyProtein. Translation follows a straightforward rule: H (P) toyN translate into H (P) toyA. Point mutations in `toy`LIFE are easy to implement: they are changes in one of the nucleotides in one of the genes in the genotype. If the sequence has a H toyN in that position, then a mutation will change it to a P toyN, and vice versa.

##### toyProteins

toyProteins correspond to the minimum energy, maximally compact folded structure of the 16 toyA chain arising from a translated toyGene. Their folded configuration is calculated through the hydrophobic-polar (HP) protein lattice model [40, 41].

We only consider maximally compact structures. That is, every toyProtein must fold on a 4 × 4 lattice, following a self-avoiding walk (SAW) on it. After accounting for symmetries —rotations and reflections—, there are only 38 SAWs on that lattice (Supplementary Figure S2).

**Supplementary Figure S2:**
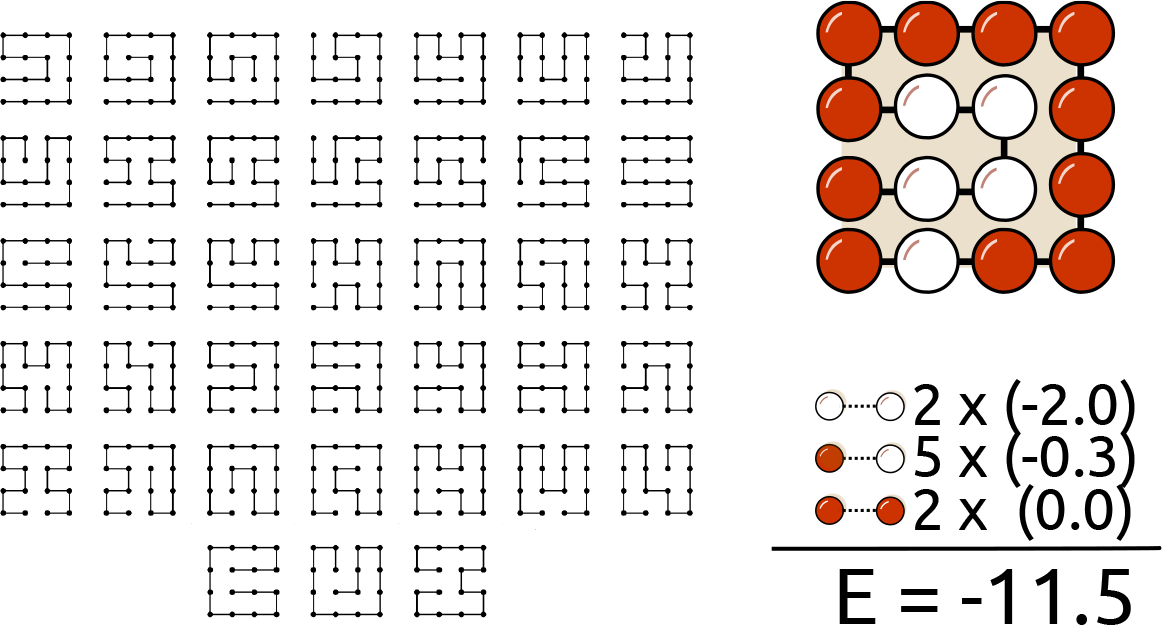
Protein folding in toyLIFE. toyProteins fold on a 4 × 4 lattice, following a self-avoiding walk (SAW). Discarding for symmetries, there are 38 SAWs (left). For each binary sequence of length 16, we fold it into every SAW and compute its folding energy, following the HP model. For instance, we fold the sequence PHPPPPPPPPPHHHHP into one of the SAWs and compute its folding energy (right). There are two HH contacts, five HP contacts and two PP contacts —we only take into account contacts between non-adjacent toyAminoacids. Summing all this contacts with their corresponding energies, we obtain a folding energy of −11.5. Repeating this process for every SAW, we obtain the minimum free structure.

The energy of a fold is the sum of all pairwise interaction energies between toyA that are not contiguous along the sequence. Pairwise interaction energies are *E*_HH_ = −2, *E*_HP_ = −0.3 and *E*_PP_ = 0, following the conditions set in [40] that *E*_PP_ > *E*_HP_ > *E*_HH_ (Supplementary Figure S2). toyProteins are identified by their folding energy and their perimeter. If there is more than one fold with the same minimum energy, we select the one with fewer H toyAminoacids in the perimeter. If still there is more than one fold fulfilling both conditions, we discard that protein by assuming that it is intrinsically disordered and thus non-functional. Note, however, that sometimes different folds yield the same folding energy and the same perimeter. In those cases, we do not discard the resulting toyProtein.

Out of 2^16^ = 65, 536 possible toyProteins, 12, 987 do not yield unique folds. We find 2, 710 different toyProteins with 379 different perimeters. Not all toyProteins are equally abundant: although every toyProtein is coded by 19.4 toyGenes on average, most of them are coded by only a few toyGenes. For instance, 1, 364 toyProteins —roughly half of them!— are coded by less than 10 toyGenes each. On the other hand, only 4 toyProteins are coded by more than 200 toyGenes each, the maximum being 235 toyGenes coding for the same toyProtein. The distribution is close to an exponential decay (Supplementary Figure S3**a**). The same happens with the perimeters, although with less skewness: each perimeter is mapped by 7.15 toyProteins on average, but the most abundant perimeters correspond to 26 toyProteins, and 100 are mapped by 1 or 2 toyProteins each (Supplementary Figure S3**b**).

Folding energies range from −18.0 to −0.6, with an average in −9.63. The distribution is unimodal, although very rugged (Supplementary Figure S3**c**). Note that folding energies are discrete, and that separations between them are not equal. For instance, there are 6 toyProteins that have a folding energy of −18.0, but the next energy level is −16.3, realised by 17 toyProteins, and yet the next level is −16.0, realised by 14 toyProteins. The mode of the distribution is −10.6, realised by 202 toyProteins.

We can also study the structure of the toyProtein network (Supplementary Figure S3**e**, **f**). The nodes of this network will be the 2, 710 toyProteins. toyProtein 1 and toyProtein 2 will be neighbors if there is a pair of toyGenes that express each toyProtein and whose sequence is equal but for one toyN. The weight of the edge between toyProtein1 and 2 will be the sum of such pairs of toyGenes. It is surprising that there are no self-loops in this network —there are no mutations connecting one toyProtein to itself. In other words, although there is a strong degeneracy in the mapping from toyGenes to toyProteins, there are no connected neutral networks. If we consider just the perimeters, however, the neutrality is somewhat recovered: out of the 379 perimeters, 224 of them have neutral neighbors. So there are many mutations that alter the folding energy of a toyProtein without changing the perimeter. In this sense, `toy`LIFE is capturing a complex detail of molecular biology: mutations appear to be neutral from one point of view —in this case, perimeter— but are rarely entirely neutral. In other words, the value of a mutation is context and environment-dependent. There are always some small changes in the molecule —in this case, folding energy— that may affect their function later down the line. Real world examples of this *cryptic* effects of mutations on molecules are everywhere [42–45]. Connections between toyProteins are scarce too: the average degree in the toyProtein network is 32.2 (with a standard deviation of 25.7), a very small number — on average, each toyProtein is connected to hardly 1% of the rest of toyProteins! (Supplementary Figure S3**e**). The maximum degree is 190. This means that mutating from one toyProtein to another is not easy in general. In terms of perimeters this is more relaxed, as the average degree in the perimeter network is 53.3 (standard deviation is 38.1), with a maximum degree of 173. On average, every perimeter is connected to 14% of the rest of perimeters: it is a small number, but it is still higher than in the toyProtein case (Supplementary Figure S3**f**).

**Supplementary Figure S3:**
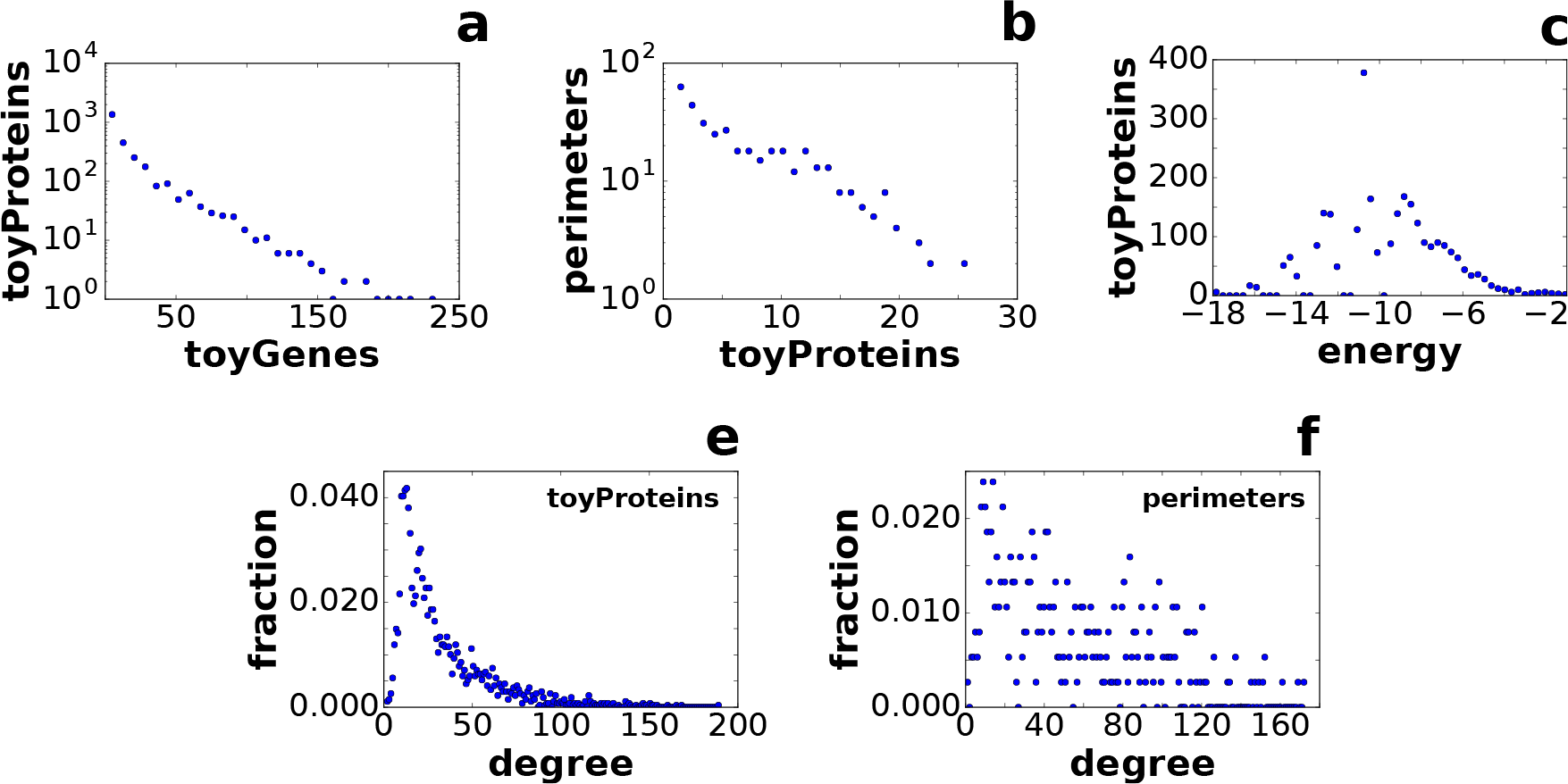
Distributions of toyProteins in toyLIFE. **(a)** Distribution of toyProtein abundances —that is, the number of toyGenes that code for them. Most toyProteins are coded by few toyGenes, but some of them are very abundant: the most abundant toyProtein is coded by 235 toyGenes. **(b)** Distribution of the perimeters associated with each toyProtein. Again, not all perimeters are equally abundant, and some of them correspond to as many as 25 toyProteins, while 100 correspond to 1 or 2 toyProteins. **(c)** Distribution of folding energies. The range of folding energies goes from −18.0 to −0.6, with a unimodal, rugged distribution. The mode is −10.6, a folding energy achieved by 202 toyProteins. **(d)** Degree distribution in the toyProtein network. Two toyProteins are connected if there are two toyGenes coding for them that have the same sequence, except for one toyN. The average degree is 32.2. **(e)** Degree distribution in the perimeter network. Two perimeters are neighbors if the toyProteins associated to them are neighbors. The average degree is 53.3.

In the `toy`LIFE universe, only the folding energy and perimeter of a toyProtein matter to characterise its interactions, so folded chains sharing these two features are indistinguishable. This is a difference with respect to the original HP model, where different inner cores defined different proteins and the composition of the perimeter was not considered as a phenotypic feature. However, subsequent versions of HP had already included additional traits [46].

The toyPolymerase (Supplementary Figure S1) is a special toyA polymer, similar to a toyProtein in many aspects, but that is not coded for by any toyGene. It has only one side, with sequence PHPH, and its folding energy is taken to be *−*11.0. We will discuss its function and place later on.

#### 1.2 Extending the HP model: interactions

toyProteins interact through any of their sides with other toyProteins, with promoters of toyGenes, and with toyMetabolites (see Supplementary Figure S4**a**). When toyProteins bind to each other, they form a toyDimer, which is the only protein aggregate considered in `toy`LIFE. The two toyProteins disappear, leaving only the toyDimer. Once formed, toyDimers can also bind to promoters or toyMetabolites through any of their sides —binding to other toyProteins or toyDimers, however, is not permitted. In all cases, the interaction energy (*E*_int_) is the sum of pairwise interactions for all HH, HP and PP pairs formed in the contact —these interactions follow the rules of the HP model as well. Bonds can be created only if the interaction energy between the two molecules *E*_int_ is lower than a threshold energy *E*_thr_ = −2.6. Note that a minimum binding energy threshold is necessary to avoid the systematic interaction of any two molecules. Low values of the threshold would lead to many possible interactions, which would increase computation times. High values would lead to very few interactions, and we would obtain a very dull model. Our choice of *E*_thr_ = −2.6 achieves a balance: the number of interactions is large enough to generate complex behaviours, as we will see later on, while at the same time keeping the universe of interactions small enough to handle computationally. If below threshold, the total energy of the resulting complex is the sum of *E*_int_ plus the folding energy of all toyProteins involved. The lower the total energy, the more stable the complex. When several toyProteins or toyDimers can bind to the same molecule, only the most stable complex is formed. Consistently with the assumptions for protein folding, when this rule does not determine univocally the result, no binding is produced.

As the length of toyMetabolites is usually longer than 4 toyS (the length of interacting toyProtein sites), several binding positions between a toyMetabolite and a toyProtein might share the same energy. In those cases we select the sites that yield the most centered interaction (Supplementary Figure S4b). If ambiguity persists, no bond is formed. Also, no more than one toyProtein / toyDimer is allowed to bind to the same toyMetabolite, even if its length would permit it. toyProteins / toyDimers bound to toyMetabolites cannot bind to promoters.

**Supplementary Figure S4:**
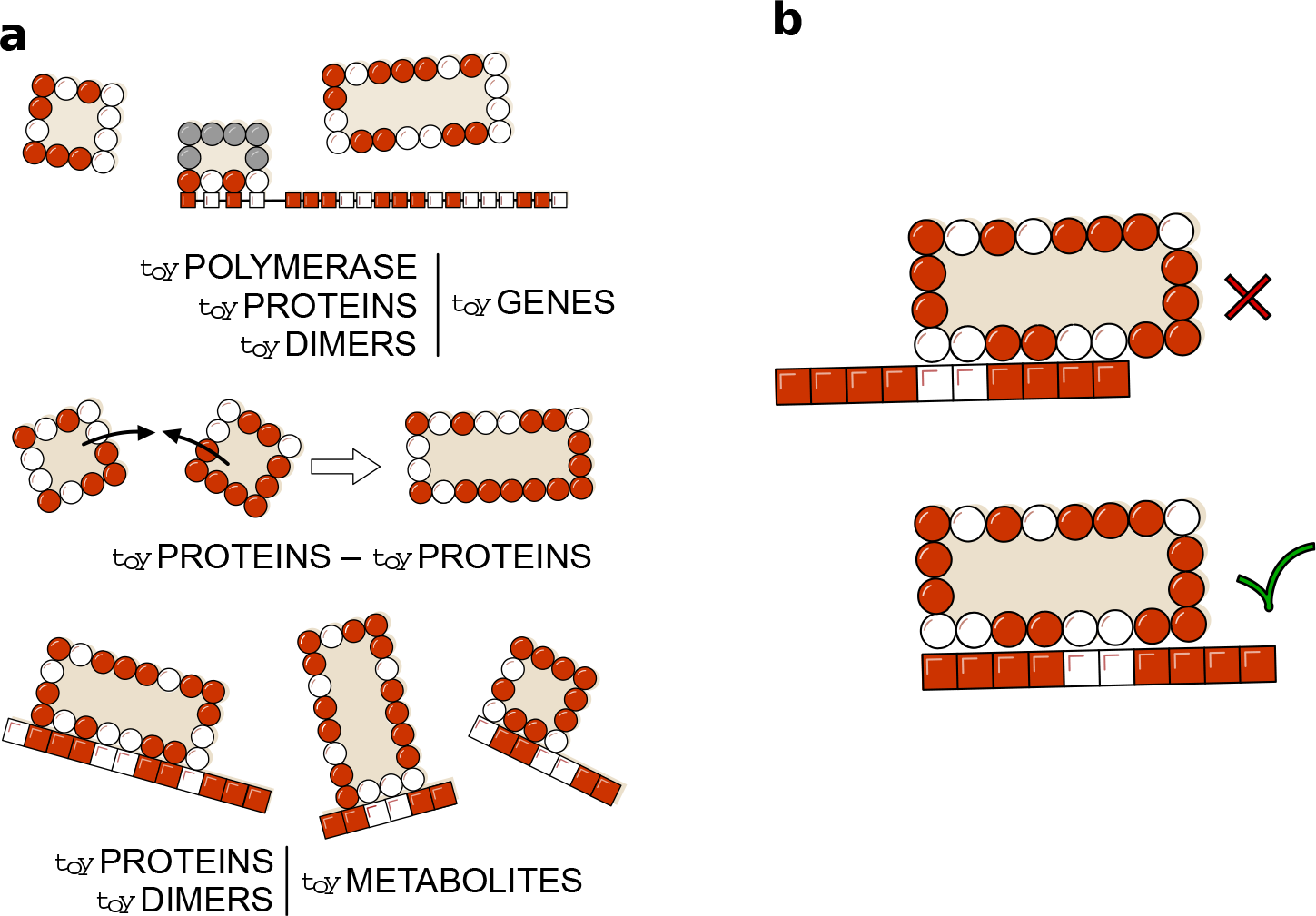
Interactions in toyLIFE. **(a)** Possible interactions between pairs of toyLIFE elements. toyGenes interact through their promoter region with toyProteins (including the toyPolymerase and toyDimers); toyProteins can bind to form toyDimers, and interact with the toyPolymerase when bound to a promoter; both toyProteins and toyDimers can bind a toyMetabolite at arbitrary regions along its sequence. **(b)** When a toyDimer or toyProtein binds to a toyMetabolite with the same energy in many places, we choose the most centered binding position. If two or more binding positions have the same energy and are equally centered, then no binding occurs.

Interaction rules in `toy`LIFE have been devised to remove any ambiguity. When more than one rule could be chosen, we opted for computational simplicity, having made sure that the general properties of the model remained unchanged. A detailed list of the specific disambiguation rules implemented in the model follows:

1. **Folding rule:** if a sequence of toyAminoacids can fold into two (or more) different configurations with the same energy and two different perimeters with the same number of H, it is considered degenerate and does not fold.
2. **One-side rule:** any interaction in which a toyProtein can bind any ligand with two (or more) different sides and the same energy is discarded.
3. **Annihilation rule:** if two (or more) toyProteins can bind a ligand with the same energy, the binding does not occur. However, if a third toyProtein can bind the ligand with greater (less stable) energy than the other two, and does so uniquely, it will bind it.
4. **Identity rule:** an exception to the Annihilation rule occurs if the competing toyProteins are the same. In this case, one of them binds the ligand and the other(s) remains free.
5. **Stoichiometric rule:** an extension of the Identity rule. If two (or more) copies of the same toyProtein / toyDimer / toyMetabolite are competing for two (or more) different ligands, there will be binding if the number of copies of the toyProtein / toyDimer / toyMetabolite equals the number of ligands. For example, say that P1 binds to P2, P3 and P4 with the same energy. Then, if P1, P2 and P3 are present, no complex will form; (b) if there are two copies of P1, dimers P1-P2 and P1-P3 will both form; but (c) if P4 is added, no complex will form. Conversely, if all ligands are copies as well, the Stoichiometry rule does not apply. For example, three copies of P1 and two copies of P2 will form two copies of dimer P1-P2, and one copy of P1 will remain free.

#### 1.3 Regulation

Expression of toyGenes occurs through the interaction with the toyPolymerase, which is a special kind of toyProtein (see Supplementary Figure S1). The toyPolymerase only has one interacting side (with sequence PHPH) and its folding energy is fixed to value −11.0: it is more stable than more than half the toyProteins. It is always present in the system. The toyPolymerase binds to promoters or to the right side of a toyProtein / toyDimer already bound to a promoter. When the toyPolymerase binds to a promoter, translation is directly activated and the corresponding toyGene is expressed (Supplementary Figure S5**a**). However, a more stable (lower energy) binding of a toyProtein or toyDimer to a promoter precludes the binding of the toyPolymerase. This inhibits the expression of the toyGene, except if the toyPolymerase binds to the right side of the toyProtein / toyDimer, in which case the toyGene can be expressed.

The minimal interaction rules that define `toy`LIFE dynamics endow toyProteins with a set of possible activities not included *a priori* in the rules of the model (see Supplementary Figure S5). For example, since the 4-toyN interacting site of the toyPolymerase cannot bind to all promoter regions —because some of these interactions have *E*_int_ > *E*_thr_—, translation mediated by a toyProtein or toyDimer binding might allow the expression of genes that would otherwise never be translated. These toyProteins thus act as activators (Supplementary Figure S5**c**). This process finds a counterpart in toyProteins that bind to promoter regions more stably than the toyPolymerase does, and therefore prevent gene expression — this happens if *E*_int(PROT)_ + *E*_PROT_ < *E*_int(POLY)_ + *E*_POLY_. They are acting as inhibitors (Supplementary Figure S5**b**). There are two additional functions that could not be foreseen and involve a larger number of molecules. A toyProtein that forms a toyDimer with an inhibitor —preventing its binding to the promoter— effectively behaves as an activator for the expression of the toyGene. However, it interacts neither with the promoter region nor with the toyPolymerase, and its activating function only shows up when the inhibitor is present. This toyProtein thus acts as a conditional activator (Supplementary Figure S5**d**). On the other hand, two toyProteins can bind together to form a toyDimer that inhibits the expression of a particular toyGene. As the presence of both toyProteins is needed to perform this function, they behave as conditional inhibitors (Supplementary Figure S5**e**). This flexible, context-dependent behavior of toyProteins is reminiscent of phenomena observed in real cells [47], and permits the construction of complex toyGene Regulatory Networks (toyGRNs).

**Supplementary Figure S5:**
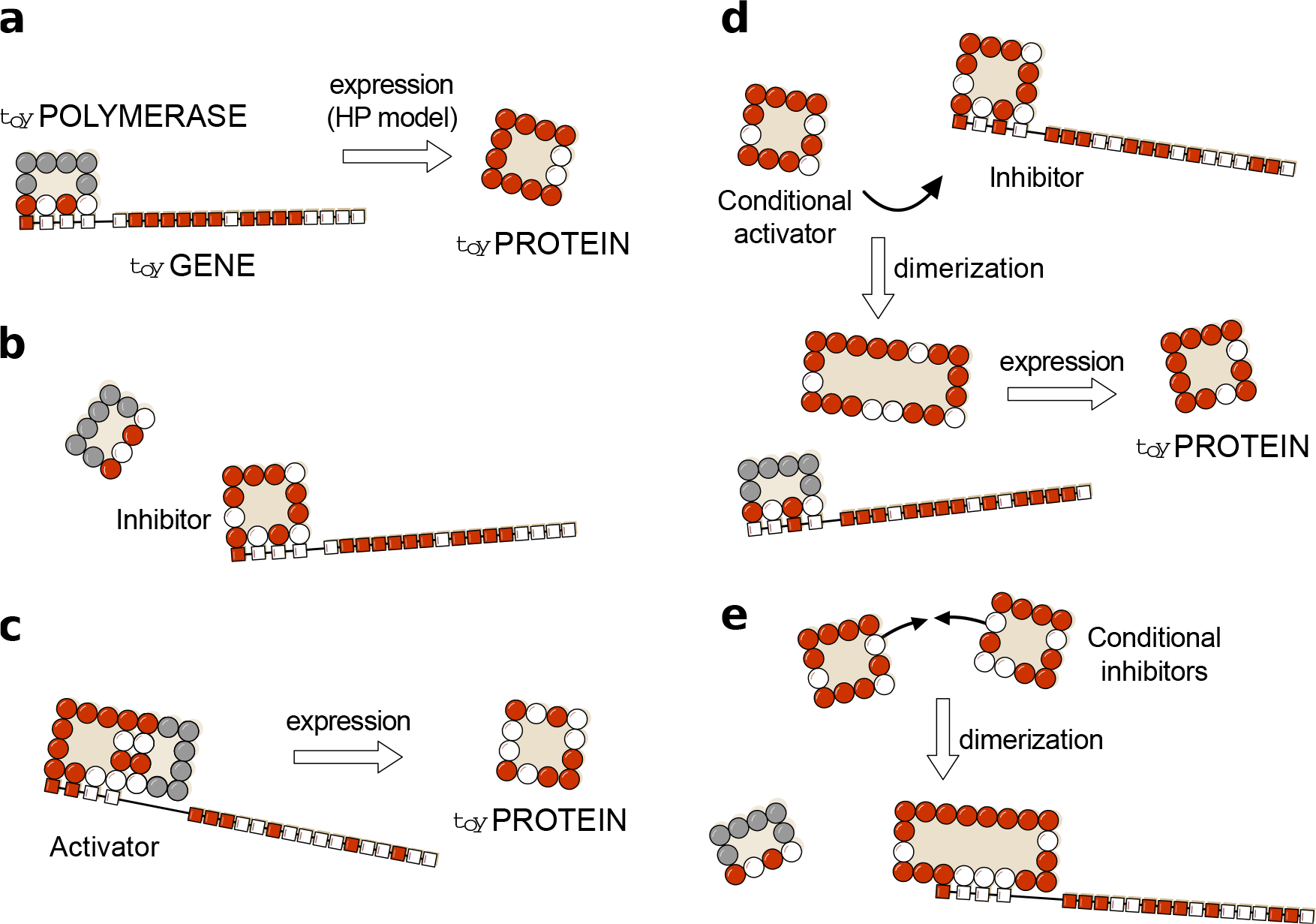
Regulatory functions in toyLIFE. **(a)** A toyGene is expressed (translated) when the toyPolymerase binds to its promoter region. The sequence of Ps and Hs of the toyProtein will be exactly the same as that of the toyGene coding region. **(b)** If a toyProtein binds to the promoter region of a toyGene with a lower energy than the toyPolymerase does, it will displace the latter, and the toyGene will not be expressed. This toyProtein acts as an *inhibitor*. **(c)** The toyPolymerase does not bind to every promoter region. Thus, not all toyGenes are expressed constitutively. However, some toyProteins will be able to bind to these promoter regions. If, once bound to the promoter, they bind to the toyPolymerase with their rightmost side, the toyGene will be expressed, and these toyProteins act as *activators*. **(d)** More complex interactions —involving more elements— appear. For example, a toyProtein that forms a toyDimer with an inhibitor —preventing it from binding to the promoter— will effectively activate the expression of the toyGene. However, it does neither interact with the promoter region nor with the toyPolymerase, and its function is carried out only when the inhibitor is present. We call this kind of toyProteins *conditional activators*. **(e)** Two toyProteins can bind together to form a toyDimer that inhibits the expression of a certain toyGene. As they need each other to perform this function, we call them *conditional inhibitors*. As the number of genes increases, this kind of complex relationships can become very intricate.

#### 1.4 Metabolism

When a toyDimer is bound to a toyMetabolite, another toyProtein can interact with this complex and break it. This reaction will take place if the toyProtein can bind to one of the subunits of the toyDimer and the resulting complex has less total energy than the toyDimer. As with the rest of interactions, the catabolic reaction will only take place if this binding is unambiguous. As a result of this reaction, the toyDimer will be broken in two: one of the pieces will be bound to the toyProtein (forming a new toyDimer), and the other one will remain free. The toyMetabolite will break accordingly: the part of it that was bound to the first subunit will stay with it, and the other part will stay with the second subunit. Note that the toyMetabolite need not be broken symmetrically: this will depend on how the toyDimer binds to it (Supplementary Figure S6).

**Supplementary Figure S6:**
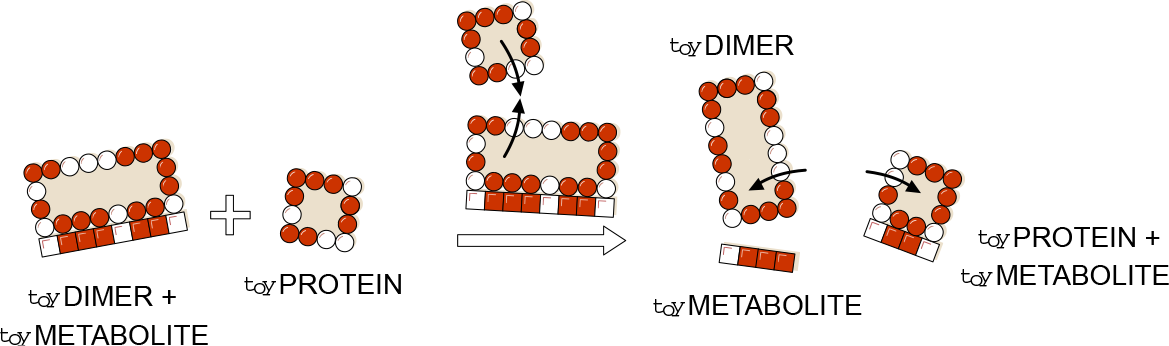
Metabolism in toyLIFE. A toyDimer is bound to a toyMetabolite when a new toyProtein comes in. If the new toyProtein binds to one of the two units of the toyDimer, forming a new toyDimer energetically more stable than the old one, the two toyProteins will unbind and break the toyMetabolite up into two pieces. We say that the toyMetabolite has been catabolised.

#### 1.5 Dynamics in toyLIFE

The dynamics of the model proceeds in discrete time steps and variable molecular concentrations are not taken into account. A step-by-step description of `toy`LIFE dynamics is summarised in Supplementary Figure S7. There is an initial set of molecules which results from the previous time step: toyProteins (including toyDimers and the toyPolymerase) and toyMetabolites, either endogenous or provided by the environment. These molecules first interact between them to form possible complexes (see Section 1.2) and are then presented to a collection of toyGenes that is kept constant along subsequent iterations. Regulation takes place, mediated by a competition for binding the promoters of toyGenes, possibly causing their activation and leading to the formation of new toyProteins. Binding to promoters is decided in sequence. Starting with any of them (the order is irrelevant), it is checked whether any of the toyProteins / toyDimers (including the toyPolymerase) available bind to the promoter —remember that complexes bound to toyMetabolites are not available for regulation—, and then whether the toyPolymerase can subsequently bind to the complex and express the accompanying coding region. If it does, the toyGene is marked as active and the toyProtein / toyDimer is released. Then a second promoter is chosen and the process repeated, until all promoters have been evaluated. toyGenes are only expressed after all of them have been marked as either active or inactive. Each expressed toyGene produces one single toyProtein molecule. There can be more units of the same toyProtein, but only if multiple copies of the same toyGene are present.

**Supplementary Figure S7:**
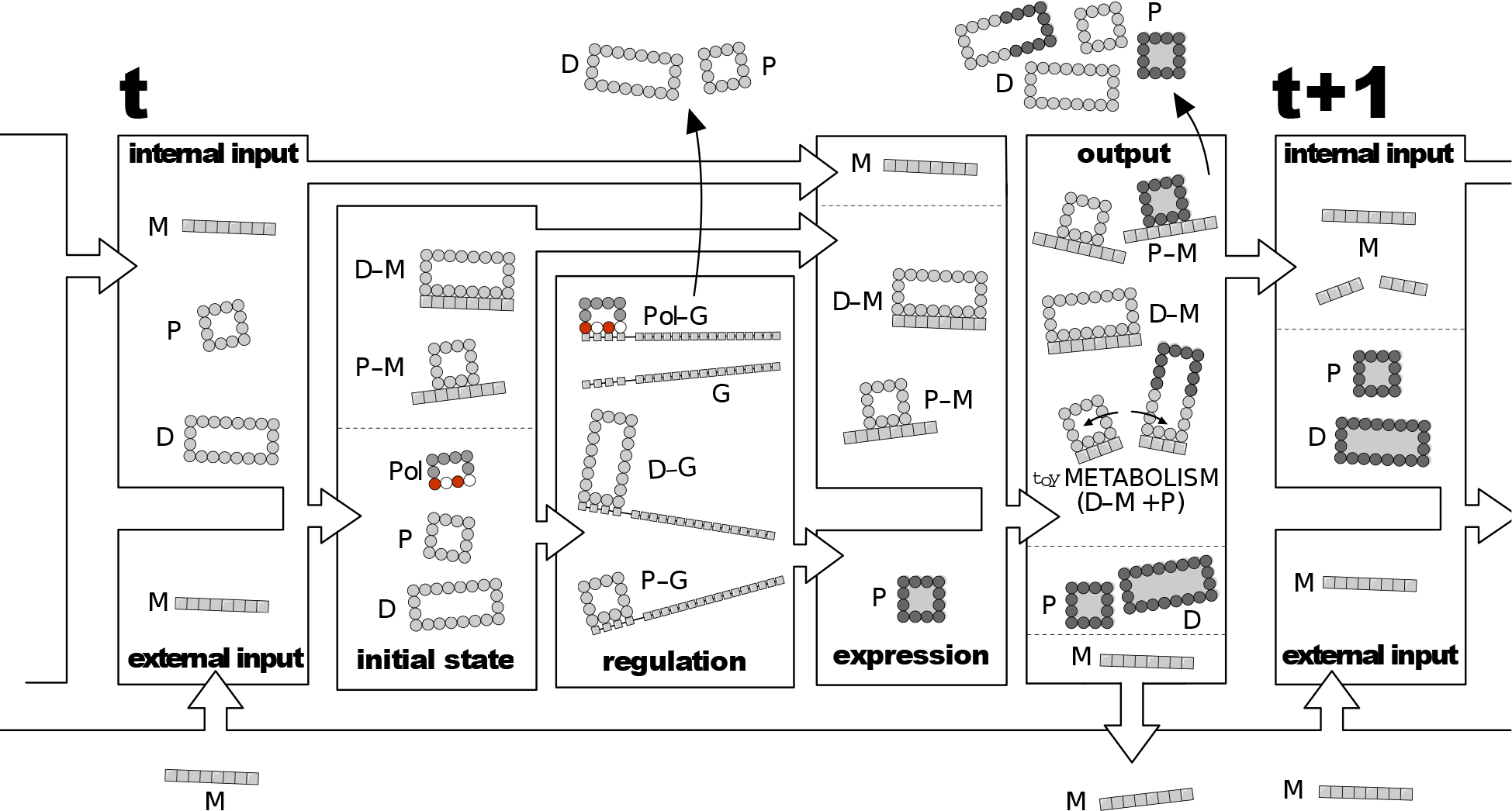
Dynamics of toyLIFE. Input molecules at time step *t* are toyProteins (P_s_) (including toyDimers (D_s_)) and toyMetabolites, either produced as output at time step *t* − 1 or environmentally supplied (all toyMetabolites denoted Ms). Ps and Ds interact with Ms to produce complexes P-M and D-M. Next, the remaining P_s_ and D_s_ and the toyPolymerase (Pol) interact with toyGenes (G) at the regulation phase. The most stable complexes with promoters are formed (Pol-G, P-G and D-G), activating or inhibiting toyGenes. P-Ms and D-Ms do not participate in regulation. P_s_ and D_s_ not in complexes are eliminated and new P_s_ (dark grey) are formed. These P_s_ interact with all molecules present and form D_s_, new P-M and D-M complexes, and catabolise old D-M complexes. At the end of this phase, all Ms not bound to P_s_ or D_s_ are returned to the environment, and all P_s_ and D_s_ in P-M and D-M complexes unbind and are degraded. The remaining molecules (Ms just released from complexes, as well as all free P_s_ and D_s_) go to the input set of time step *t* + 1.

toyProteins / toyDimers not bound to any toyMetabolite are eliminated in this phase. Thus, only the newly expressed toyProteins and the complexes involving toyMetabolites in the input set remain. All these molecules interact yet again, and here is where catabolism can occur. Catabolism happens when, once a toyMetabolite-toyDimer complex is formed, an additional toyProtein binds to one of the units of the toyDimer with an energy that is lower than that of the initial toyDimer. In this case, the latter disassembles in favor of the new toyDimer, and in the process the toyMetabolite is broken, as already mentioned in Section 1.4 and Supplementary Figure S6. The two pieces of the broken toyMetabolites will contribute to the input set at the next time step, as will free toyProteins / toyDimers. However, toyProteins / toyDimers bound to toyMetabolites disappear in this phase —they are degraded—, and only the toyMetabolites are kept as input to the next time step. Unbound toyMetabolites are returned to the environment. This way, the interaction with the environment happens twice in each time step: at the beginning and at the end of the cycle.

## Supplementary Figures

**Supplementary Figure S8:**
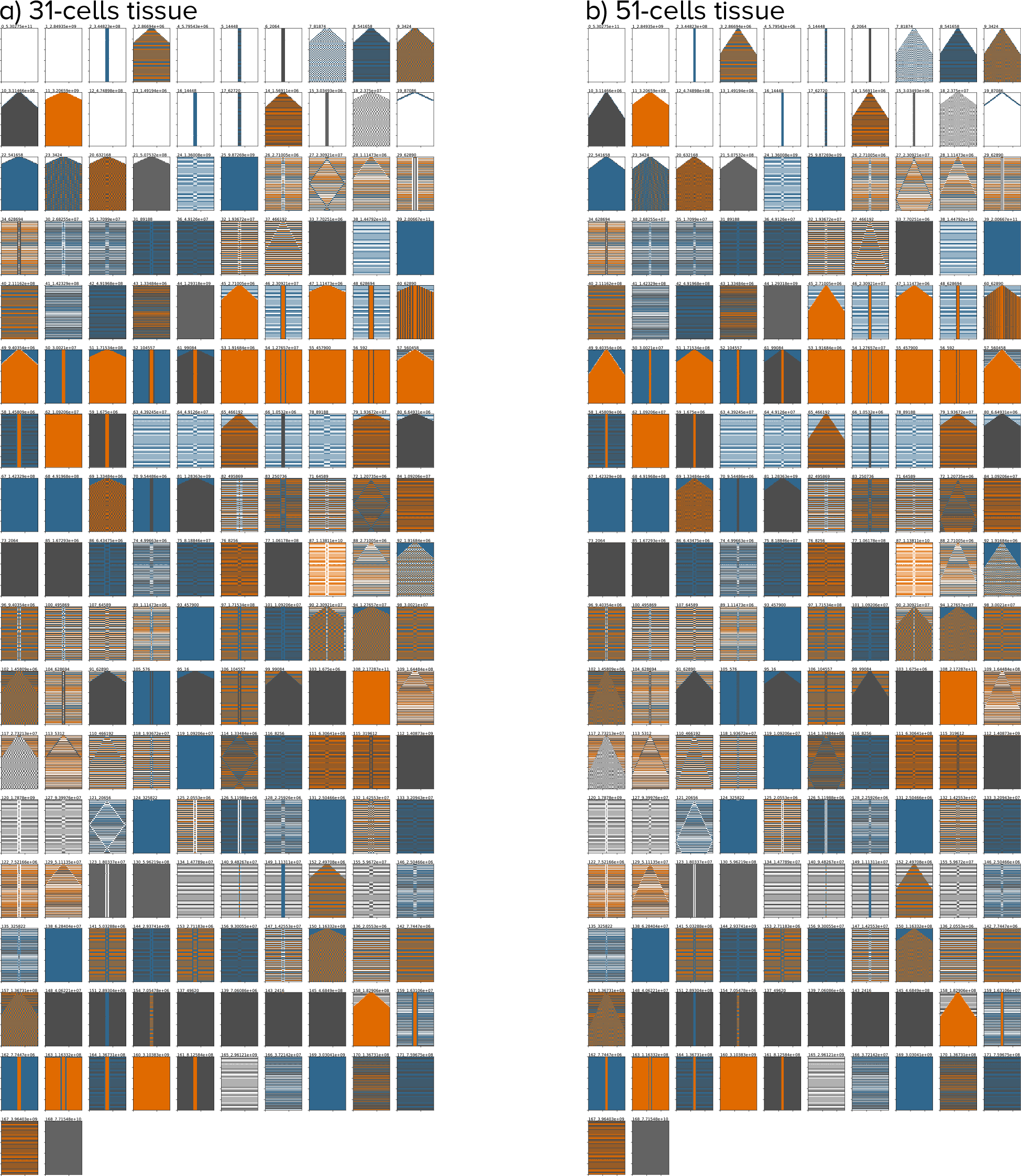
The same patterns are observed as we increase tissue size. **a)** All patterns generated by `toy`LIFE genotypes when the tissue size is set to be 31 cells. The two numbers above each pattern represent the pattern’s id and its abundance in genotype space. **b)** Same but with 51-cell tissues. The patterns are exactly the same, with the same abundances in genotype space.

**Supplementary Figure S9:**
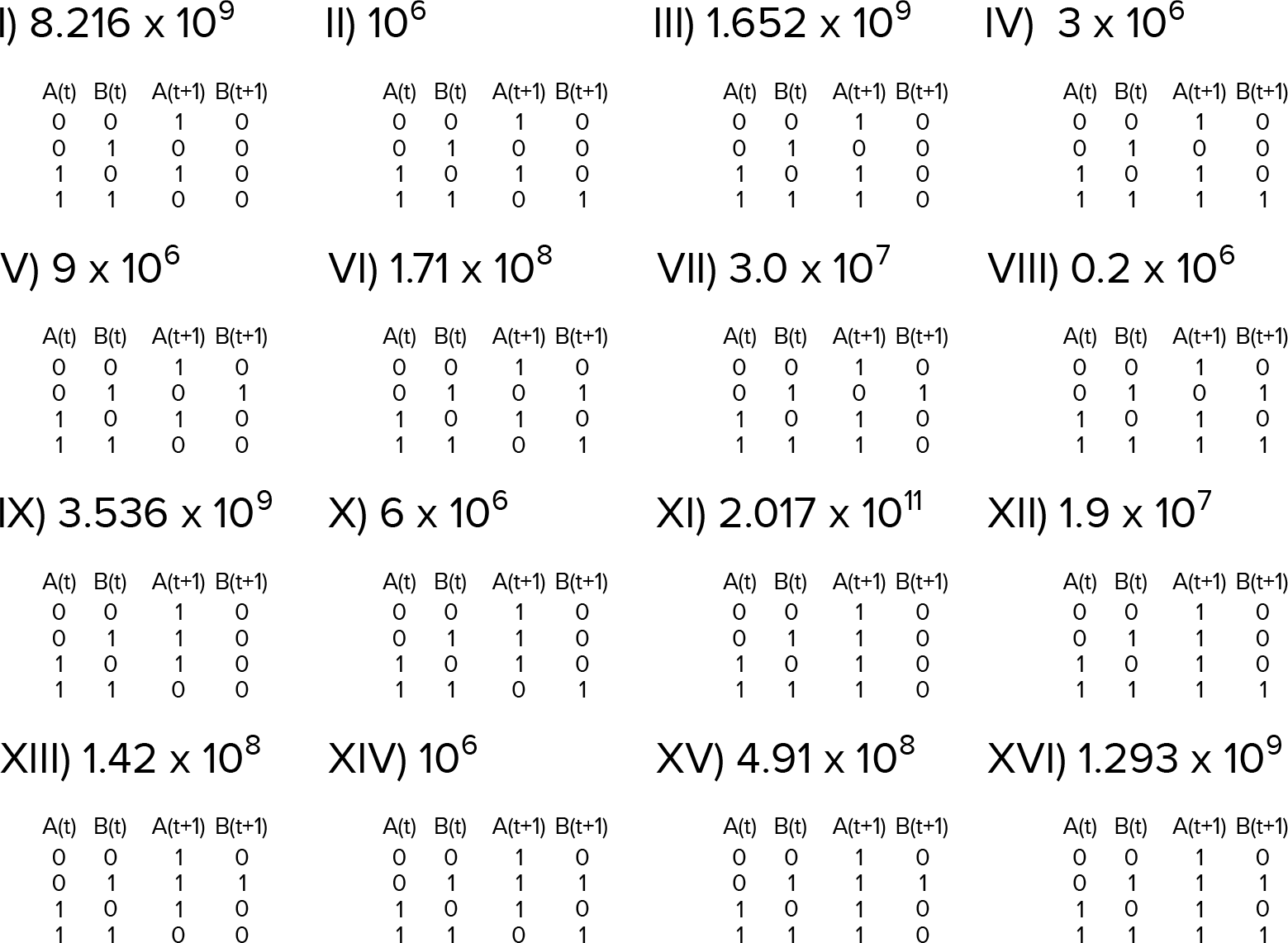
There are sixteen GRNs that generate the pattern in Figure 3b. Truth tables for all GRNs that generate the desired pattern. The number next to the label represents how many genotypes (binary sequences of length 40) are mapped into that particular GRN. Notice the wide variation in abundances.

**Supplementary Figure S10:**
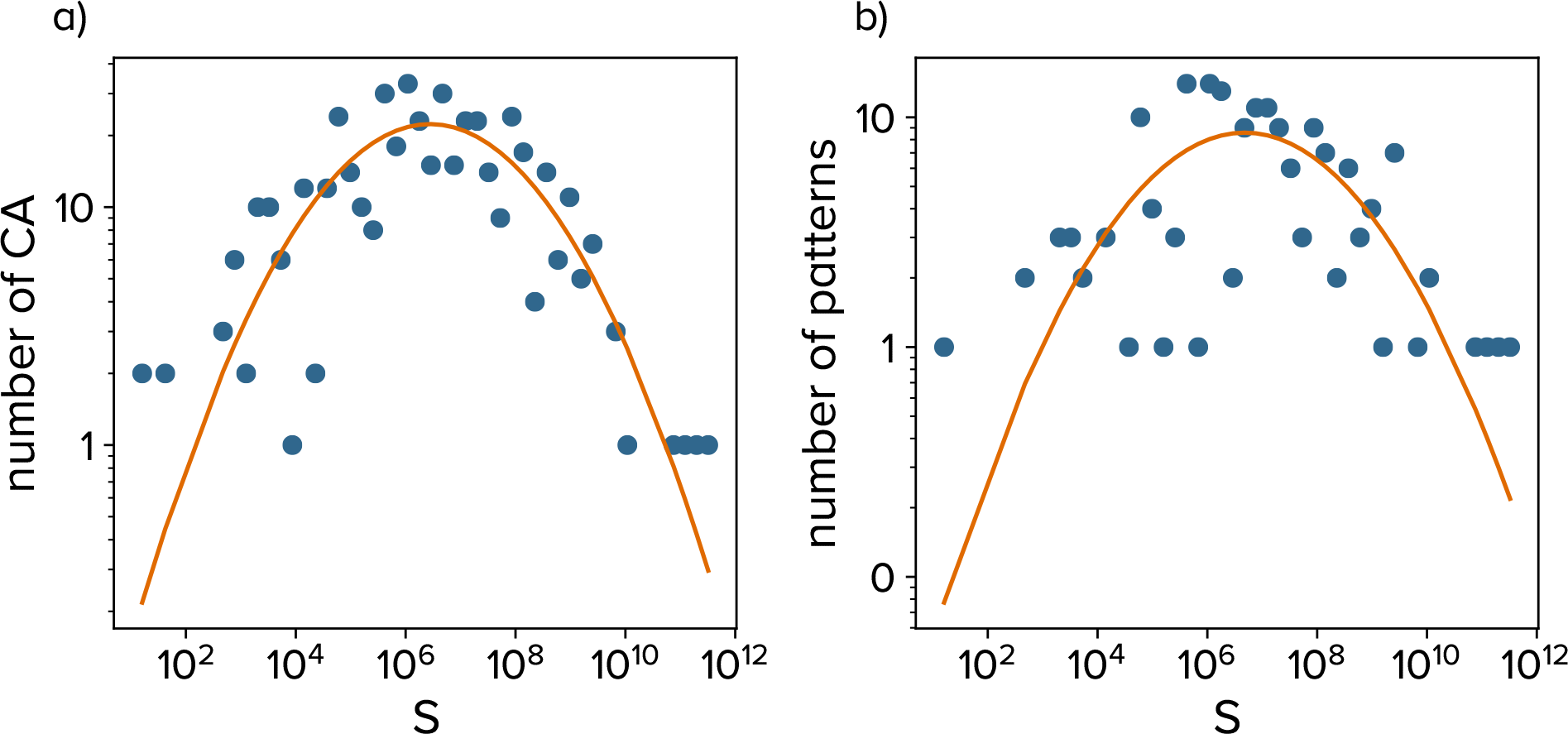
Phenotypic bias is observed in the distribution of abundances at all phenotypic levels. **(a)** The distribution of abundances of cellular automata (CA) follows a log-normal law, just like the distribution of GRNs (*R*^2^ = 0.64). **(b)** Likewise, the distribution of abundances of patterns can also be fitted by a log-normal distribution, although the fit is rather noisy (*R*^2^ = 0.41), given that we only have 172 patterns to fit.

**Supplementary Figure S11:**
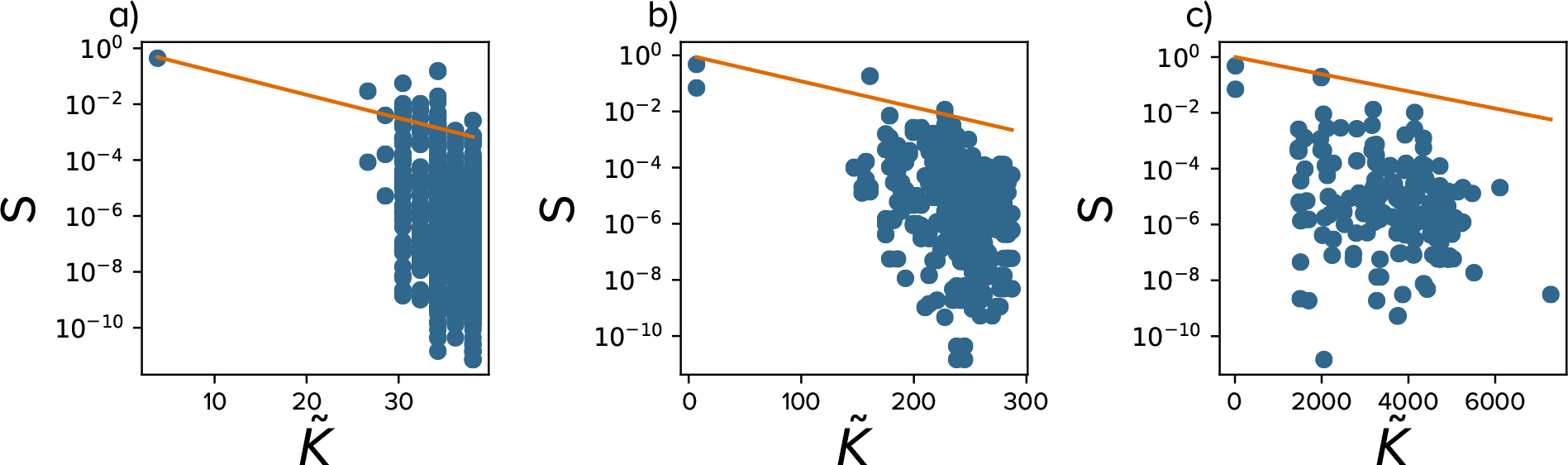
Simple phenotypes are more common in genotype space. We approximated the algorithmic complexity 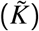 of GRNs (**a**), cellular automata (**b**) and patterns (**c**) following the work by Dingle *et al.* [29] (Methods), and plotted them against phenotype abundance (*S*). The disparity in lengths between the string representation of different phenotypic levels explains the difference in magnitude in the values of 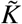. Dingle *et al.* conjecture that many input-output maps have the property that simple outputs (as measured by their algorithmic complexity) should be mapped by more inputs. In our case, this would mean that simple phenotypes are more abundant in genotype space. This figure confirms this prediction for our three phenotypic levels. Lines represent the upper bound computed in [29], 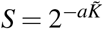, 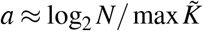, where *N* is the number of phenotypes and the maximal 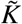 is computed over all possible phenotypes (which is straightforward in our case as we know the complete maps). GRNs and cellular automata do not always lie below the upper bound. This could be explained because the results obtained by Dingle *et al.* rely on asymptotic approximations with long strings, but the strings coding these two phenotypic levels are not very long, so asymptotic approximations may fail.

**Supplementary Figure S12:**
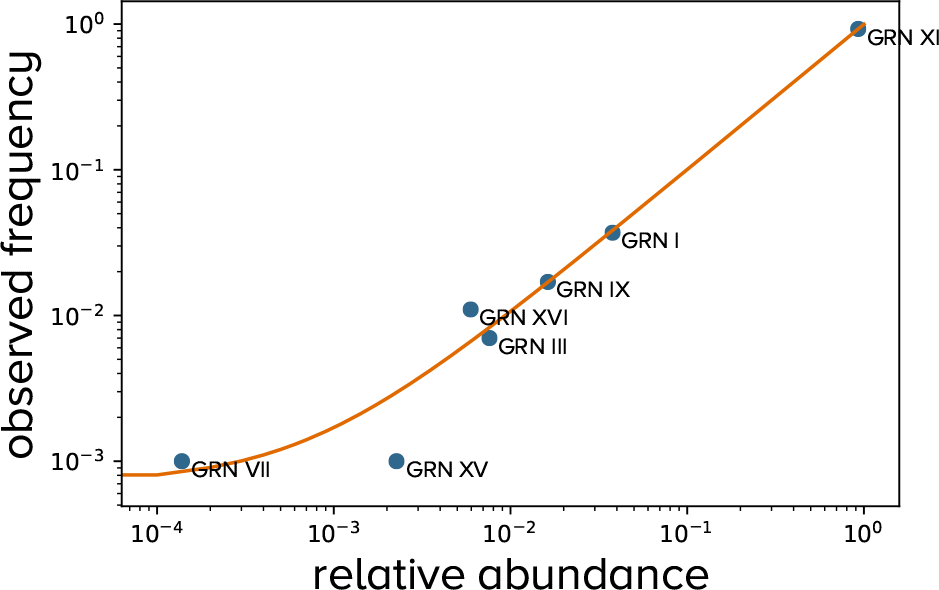
Equally fit GRNs appear as the endpoint of evolutionary simulations in proportion to their relative abundance in genotype space. Although all sixteen GRNs are equally fit (see main text), evolutionary simulations in which populations undergo Wright-Fisher dynamics do not find every GRN with equal probability. On the contrary, those GRNs that are more abundant in genotype space appear more frequently as an endpoint of our simulations, in agreement with Refs. [15, 30]. In fact, the fraction of times a given GRN is the endpoint of the simulations is almost exactly its abundance in genotype space relative to that of all sixteen GRNs. Linear fit is approximately *y* = *x* (*R*^2^ ≈ 1.0).

